# Structural insight into rabies virus neutralization revealed by an engineered antibody scaffold

**DOI:** 10.1101/2023.10.30.564668

**Authors:** Ashwini Kedari, Rommel Iheozor-Ejiofor, Lev Levanov, Kalle Saksela, Olli Vapalahti, Ilona Rissanen

## Abstract

Host-cell entry of the highly pathogenic rabies virus (RABV) is mediated by trimeric glycoprotein (G) spikes, which also represent the primary target for the humoral immune response. RABV-G displays several antigenic sites targeted by neutralizing antibodies, including monoclonal antibodies (mAbs) which have been proposed as quality-controlled alternatives to traditional polyclonal rabies immunoglobulin treatment. In this study, we determine the epitope of a potently neutralizing human anti-rabies mAb, CR57, which we engineered into a diabody to facilitate crystallization. We report the crystal structure of the CR57 diabody alone at 2.38 Å resolution, and in complex with RABV-G domain III at 3.15 Å resolution. CR57 is demonstrated to bind RABV through a predominantly hydrophobic interface, with essential interactions targeting a conserved six-residue peptide sequence ’KLCGVL’ on the RABV-G. Further, our structural analysis suggests that CR57 sterically hinders receptor recognition and the fusogenic transitions of the spike protein that are required for host-cell entry. Altogether, this investigation provides a structural perspective on rabies inhibition by a potent antibody and delineates a functionally significant region in the spike. This understanding could pave the way for the development of prophylactic antibodies and other therapeutic strategies.

**Author summary:** Rabies virus (RABV) and many other lyssaviruses possess the ability to invade the central nervous system, leading to fatal encephalitis in mammals. Initiation of the infectious cycle depends on host cell recognition and entry, which is mediated by viral surface glycoprotein (G) spikes and can be inhibited by spike-targeting neutralizing antibodies. In our study, we elucidated the crystal structure of an antigenic domain from RABV-G in complex with a diabody derived from the potently neutralizing antibody CR57. This investigation revealed the molecular interactions by which CR57 binds to RABV-G and outlined a site of vulnerability comprising a conserved peptide in RABV-G domain III, where antibody binding is likely to inhibit RABV by obstructing host cell entry. Insights into the binding modalities of antibodies like CR57 deepen our understanding of how RABV and other lyssaviruses are neutralized, aiding the development of potential therapeutics. Furthermore, our study showcases the utility of engineering antibodies into diabodies to obtain crystal structures of antibody-antigen complexes.

## Introduction

Rabies is a deadly zoonotic disease caused by the rabies virus (RABV), which belongs to the *Lyssavirus* genus within the *Rhabdoviridae* family of (-)ssRNA viruses [1–3]. Although effective vaccines exist, this disease claims approximately 60000 lives annually [4, 5]. The progression of rabies disease can be prevented by post-exposure prophylaxis (PEP) treatment, which for previously unvaccinated persons includes immediate administration of a four-dose vaccine regimen and polyclonal rabies immunoglobulin (RIG) [6, 7]. Recently, there has been a growing interest in employing recombinant anti-rabies monoclonal antibodies (mAbs) as a safer and more quality-controlled alternative to traditional serum-derived polyclonal human or equine RIG in PEP [8]. Two such treatments, Rabishield, comprised of a single mAb (titled 17C7 or RAB1), and Twinrab, which is a cocktail of mAbs docaravimab and miromavimab (also known as 62-71-3 and M777), have received licensing for clinical use in India [9–12]. While anti-rabies antibodies represent an emerging and promising category of therapeutics, our current understanding of how RABV is targeted by the neutralizing immune response remains limited. Here, we address this paucity of knowledge by studying an antigenic domain of the RABV glycoprotein (RABV-G) spike.

The bullet-shaped RABV nucleocapsid is enclosed by a lipid bilayer envelope that displays trimeric G spikes, which mediate host cell entry and receptor recognition [2, 3, 13], containing an ectodomain that displays the classical rhabdoviral class III fusion protein fold [13–18]. The ectodomain is subdivided into four distinct domains: the central domain, which encompasses domains I and II, domain III (referred to as the pleckstrin homology domain), and domain IV (known as the fusion domain) [15, 16, 18]. RABV-G initiates infection by binding to host-cell receptors, followed by viral uptake by endocytosis. Upon acidification within the endosome below pH 6.5, G spikes undergo conformational changes, driving the fusion between the viral and host membranes [19, 20]. This conformational transition of the spike from a prefusion state (at neutral pH) to a postfusion state (at low pH) is facilitated by hinge regions located between domains II and III, and domains III and IV [14, 15, 21]. While the structures of the intermediary steps between pre- and postfusion states remain unknown, recent insights into the RABV-G prefusion spike in complex with neutralizing antibodies are beginning to highlight key antigenic regions on RABV-G [15, 16]. Further, there exists an extensive array of neutralizing antibodies that have yet to be structurally characterized [22, 23] and which comprise prime targets for studies aimed at identifying functionally critical spike regions.

The human mAb 57 was initially isolated from individuals vaccinated with a human diploid cell rabies vaccine, and it has demonstrated the capability to neutralize multiple RABV strains, including CVS-11, ERA, PM, and several street strains [24]. Subsequently, mAb 57 underwent a series of re-designations. It was first renamed as SO57 [25], and later retitled as CR57 in a study aimed at developing a broadly neutralizing anti-rabies mAb cocktail known as CL184 [22]. Additionally, a cocktail comprising CR57 and the mAbs RV08 and RV3A5 demonstrated efficacy in neutralizing eleven RABV strains, promoting a 100% survival rate in Syrian hamsters against a lethal RABV challenge [23, 26, 27]. Depending on the rabies strain in question, the IC50 (half-maximal inhibitory concentration) of mAb CR57 was found to range from 47 pM to 4.3 nM [28].

In this study, we elucidated the structural basis of CR57-mediated RABV neutralization using X-ray crystallography to examine RABV-G in a complex with CR57, which was engineered into a diabody to enable crystallization. We report two crystal structures: i) CR57 diabody at 2.38 Å resolution, and ii) CR57 diabody complexed with RABV-G domain III at 3.15 Å resolution. The structures resolve the interactions that mediate CR57 binding, predominantly targeting a six amino-acid peptide located within domain III. Our structure-based analysis indicates that CR57 binding is incompatible with the fusogenic transitions of the spike protein and predicted to sterically interfere with receptor binding.

## Results

### Structure and neutralizing activity of a monospecific diabody based on rabies-neutralizing mAb CR57

For the structural characterization of antibody-antigen complexes using crystallography, it is crucial to reduce the flexibility inherent to full-length mAbs. Conventionally, this is achieved by using more compact antibody fragments, such as the antigen-binding fragment (Fab) or a single chain Fv (scFv), as substitutes for the full-length mAb during crystallization [29]. In our study, while we generated a Fab from CR57, we were unable to obtain well-diffracting crystals of the Fab in complex with target antigen, RABV-G domain III. We then chose to engineer CR57 into a monospecific diabody [30]. Monospecific diabodies are composed of the heavy chain variable (V_H_) and light chain variable (V_L_) Fv-regions of an antibody fused together in a single chain, which oligomerize to form homodimers (Fig 1A). Using this approach, diffracting crystals were obtained in 2 days for the apo-diabody and in 14 days for the diabody in complex with RABV-G domain III.

**Fig 1.**
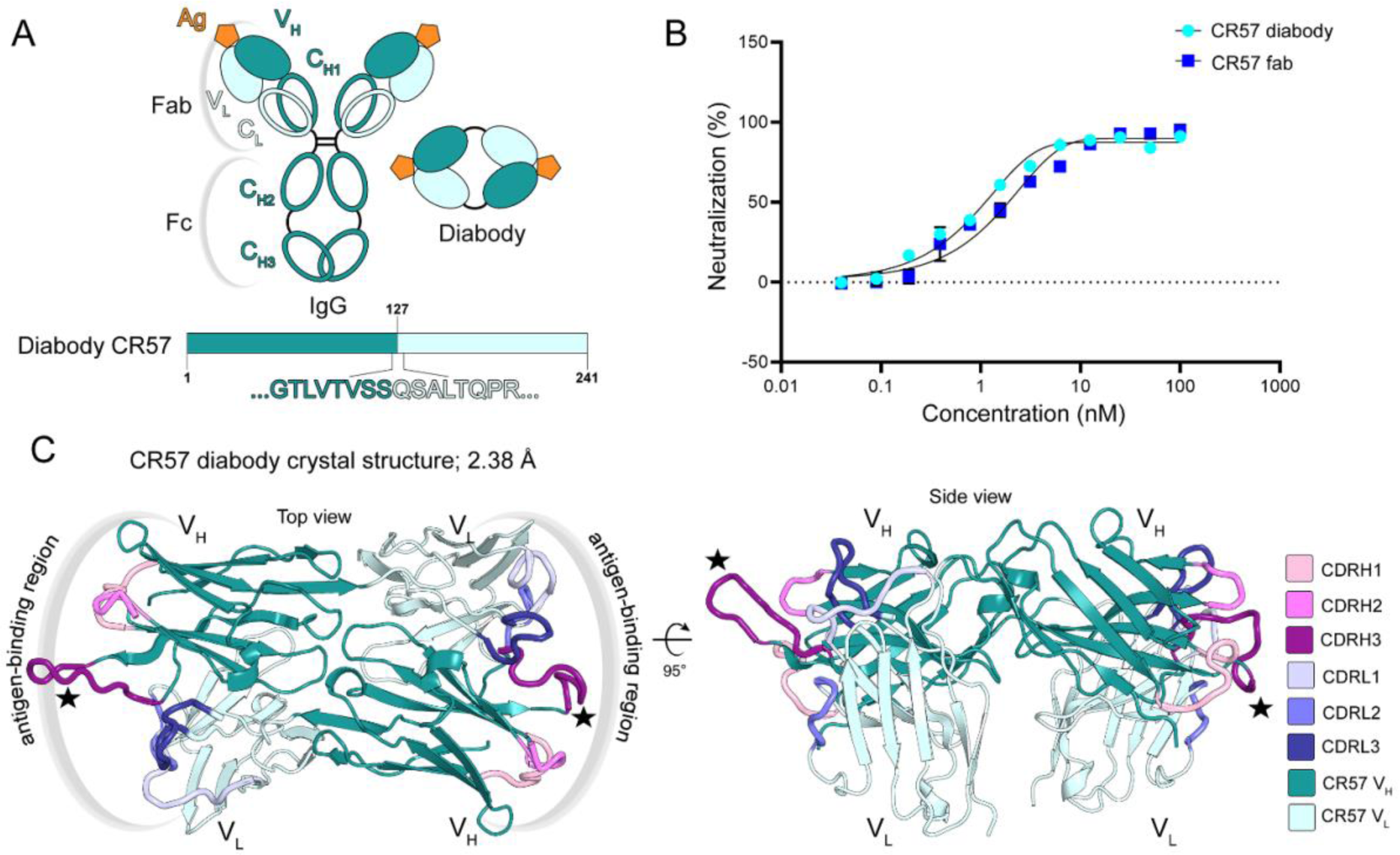
The structure of CR57 diabody and comparative neutralization properties of CR57 diabody and Fab. (A) Schematic representation of the design of CR57 diabody. Antigen (Ag) binding sites are denoted. (B) Neutralizing activity of the CR57 diabody and Fab against recombinant VSV-ΔG particles pseudotyped with RABV-G. The neutralization assay was carried out in triplicate, and error bars represent the standard deviation of three replicate measurements. (C) Crystal structure of the diabody resolved to a 2.38Å resolution. The heavy chain (V_H_) variable domain and the lambda light chain (V_L_) CDRs are shown in shades of pink and blue respectively. Conformationally variable CDRH3 is highlighted with a star.

In order to confirm that the diabody retains a functionally native antigen-binding interface, we carried out a neutralization assay utilizing recombinant VSV-ΔG particles pseudotyped with RABV-G and treated with either CR57 diabody or CR57 Fab. Both the diabody and Fab exhibited comparable neutralization capabilities, with an IC50 value of approximately 1 nM in our pseudovirus system (Fig 1B), indicating that both derivatives of the antibody share similar interactions with the antigen.

The crystal structure of the apo-CR57 diabody was resolved through X-ray crystallography to a resolution of 2.38 Å (Fig 1C, Table S1). Within the asymmetric unit (ASU), a single diabody is observed, with its two Fv-pairs oriented at an angle of 150° relative to each other. While all complementarity-determining regions (CDR) of the antibody are defined in the structure, variations in CDRH3 conformation are observed, attributable to the flexibility of the CDR loop in the absence of an antigen. Furthermore, the CR57 diabody structure displays conformational differences from the previously reported disulfide bond-stabilized diabody structures (eg. PDB 5GRX [30]) as well as from the first reported monospecific diabody structure (PDB 1LMK [31]) (Fig S1). This variability in diabody structures is due to a rotational shift of the two Fv pairs relative to each other, likely allowed by the greater flexibility of the linker regions in the diabodies PDB 1LMK and 5GRX which incorporate a five residue ‘GGGGS’ linker between V_H_ and V_L_ regions [31, 32]. In order to reduce flexibility for crystallization, no extraneous linker residues beyond the native sequence of V_H_ and V_L_ were introduced in our CR57 diabody design (Fig 1A). However, structural analysis revealed that the terminal residue S127 of the V_H_ and the initial residues Q128 and S129 of V_L_ effectively function as a linker in this construct. In our crystal structure, these residues form a bridge between the heavy and light chain domains but do not participate in the domains themselves. Additionally, when compared to native Fab models, the placement of these residues differs.

### Crystal structure of CR57 diabody in complex with RABV-G domain III reveals the CR57 epitope

To elucidate the molecular basis of RABV neutralization by mAb CR57, we resolved the crystal structure of the CR57 diabody in complex with domain III from RABV-G to 3.15 Å resolution (Fig 2, Table S1). A single complex comprised of the CR57 diabody bound to two domain III proteins was observed in the ASU. The fold of RABV-G domain III, bound to the diabody, aligns well with a previously published domain III structure (1.26 Å RMSD over 81 C-alpha atoms; PDB 7U9G, Fig S2 panel A) [33]. The CR57 epitope comprises ∼810 Å^2^ of buried surface area in domain III, and the antibody regions mediating binding are CDRH2, CDRH3, and CDRL3, which form an interface with domain III predominated by hydrophobic interactions and additionally reinforced by three hydrogen bonds (Fig 2B and Fig S3). Key interacting residues comprise Y108, Y109, and F110 in CDRH3, which form a hydrophobic interface with domain III peptide ‘KLCGVL’ (residues 226-231). This observation aligns with a previous study in which a PEPSCAN-ELISA suggested that this peptide is critical for CR57 binding [23]. Beyond the peptide ‘KLCGVL’, CDRH2 residues (I52, F55, and T57) and CDRL3 residues (G222, D223, and Y224) establish hydrophobic interactions with domain III residues M44, N194 and W251, P253, respectively (Fig S3). We also noted that the CDRL1 region presents variable conformations across the two Fv domains in the diabody. To discover if this variation might arise from potential interactions targeted at a spike region outside domain III, we superposed our diabody−domain III complex structure on the trimeric prefusion RABV-G spike (PDB 7U9G [15]). The superposition indicates that CDRL1 may engage a region in domain IV in the context of the trimeric prefusion spike (Fig S2 panel B).

**Fig 2.**
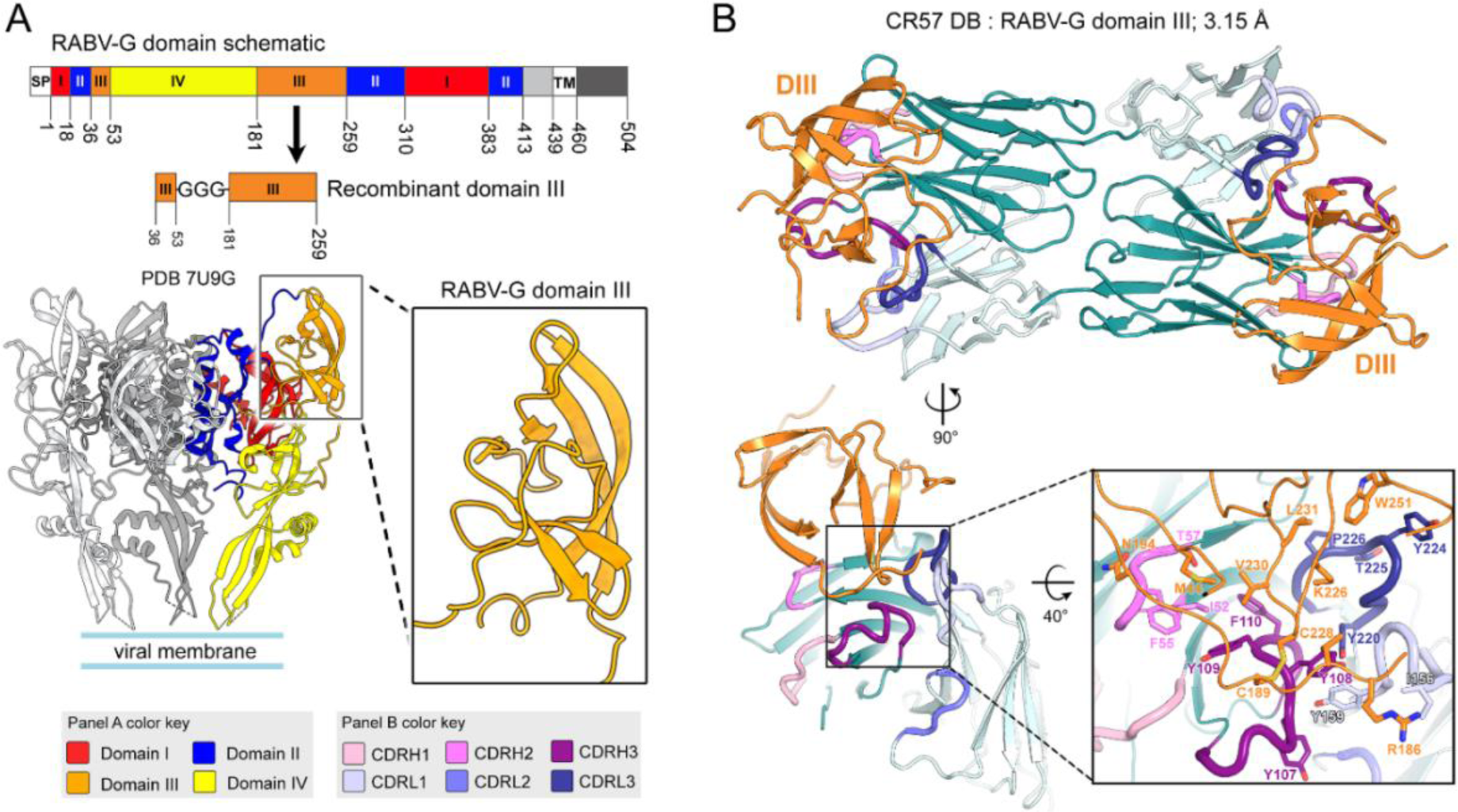
Crystal structure of the CR57 diabody in complex with RABV-G domain III. (A) Domain schematic of RABV-G shows the organization of the glycoprotein, including the signal peptide (SP), domains I-IV (colored red, blue, orange, and yellow, respectively), ectodomain C-terminus (grey), transmembrane domain (TM), and intraviral domain (black). The schematic was produced using the DOG software [60]. Composition of the domain III construct used in this study is presented alongside an illustration of domain III organization in the context of the trimeric prefusion RABV-G spike. (B) Crystal structure of the CR57 diabody in complex with RABV-G domain III, resolved at 3.15Å resolution. The lower panel includes a magnified view of the CR57 diabody−RABV-G domain III interface, where key interacting residues are labelled and shown as sticks with oxygen atoms in red and nitrogen atoms in blue.

### Analysis of antibody binding sites on RABV-G identifies crucial regions targeted for neutralization

To map sites of vulnerability to neutralization on RABV-G, we analyzed mAb epitopes on the spike (Fig 3). To date, five structures of neutralizing antibodies in complex with RABV-G have been reported, of which mAb 17C7 [16], mAb 523-11 [18], and mAb RVA122 [15] target epitopes located in domains I and II, whereas mAb RVC20 [33] and mAb 1112-1 target domain III [16]. In addition to these recent structural insights, earlier competition ELISA assays have outlined six antigenic sites (sites i-iv, minor site ‘a’ and G5) on RABV-G [22–24, 34–36] which are located in domains I to III as described in Table S2.

**Fig 3.**
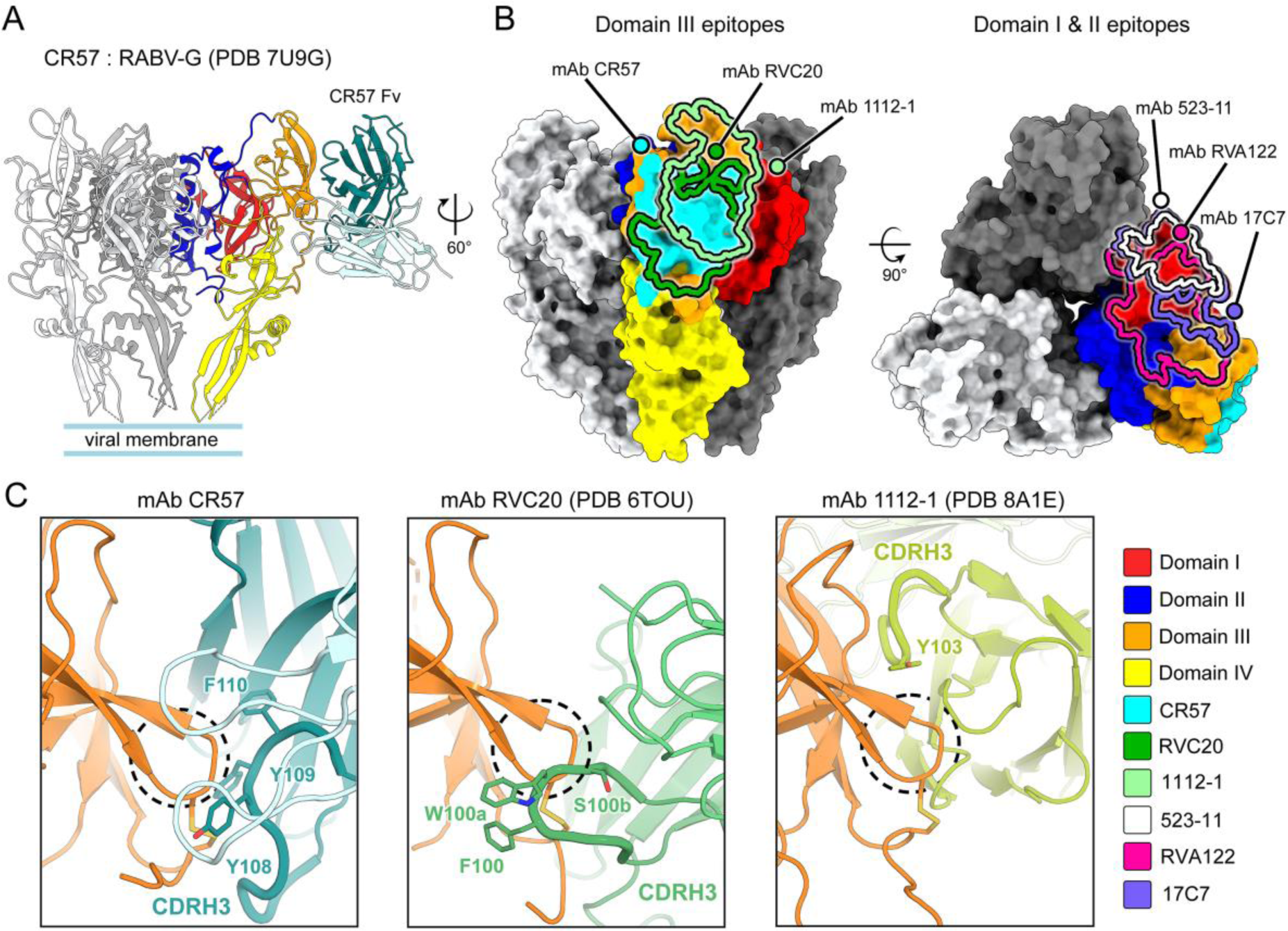
Comparative analysis of neutralizing antibody epitopes on the RABV-G spike. (A) Superposition of CR57 diabody−RABV-G domain III complex structure (for clarity, only a single CR57 Fv is shown) with a previously reported prefusion RABV-G spike (PDB: 7U9G [15]). (B) Surface representation of the spike (side and top views), displaying the epitope footprints of antibodies targeting domain III, which include CR57 (delineated in cyan), RVC20 (dark green), and 1112-1 (light green), and epitopes of antibodies targeting domains I and II, which include 523-11 (white), 17C7 (purple), and RVA122 (pink). (C) A close-up at the key interacting region between domain III and mAbs CR57; the same region is also shown for mAbs RVC20 and 1112-1 interfaces. RABV-G domain III is shown in orange, with the peptide ’KLCGVL’ highlighted by a dashed circle. CDRH3 is shown as a bold loop and residues interacting with the ’KLCGVL’ peptide are labelled and shown as sticks with oxygen atoms in red and nitrogen atoms in blue.

While epitopes of domain I and II targeting antibodies are distal from the epitope of mAb CR57, the epitopes of domain III-targeting antibodies show some overlap (Fig 3B) as expected given the relatively small accessible surface of domain III. A closer look at the residues comprising each interface reveals similarity between CR57 and RVC20, while the binding mode of mAb 1112-1 is distinct from both (Fig 3C). The conserved peptide ’KLCGVL’ (residues 226-231 in domain III; also known as RABV-G antigenic site i), is targeted by both mAb RVC20 and CR57, with key interactions mediated by aromatic residues in CDRH3 that form a hydrophobic binding pocket centered around the domain III residue C228 (Fig 3C). However, the CR57 interface is more strongly based on hydrophobic interactions, particularly with residues C228-V230 forming a β-turn, and includes an interface between CDRL3 and the C-terminal region of domain III (Fig S3) that is not targeted by RVC20.

Notably, the conserved ’KLCGVL’ peptide has been identified as a target for antibodies capable of cross-neutralizing multiple strains of RABV and related lyssaviruses [22, 23, 28]. Core residues ‘KLCGV’ of the peptide represent a conserved site, especially within lyssavirus phylogroup I, where these residues are either fully conserved or only exhibit variation with chemically similar residues (K/R, V/I) [33]. This provides a rationale for the cross-neutralization potency of antibodies targeting this region, such as mAbs CR57, RVC20, and RVC3 [28]. While the specific epitope recognized by mAb RVC3 has yet to be determined, the similarities in the interfaces of mAb CR57 and RVC20 hint that RVC3, along with other antibodies targeting antigenic site i (listed in Table S2), might employ comparable binding strategies, including a hydrophobic interface between aromatic CDRH3 residues and peptide ‘KLCGVL’.

The epitopes of neutralizing antibodies may outline functionally significant regions within the rabies spike. We note that in line with the abrogation of receptor binding as one of the mechanisms of virus neutralization, two regions implicated for receptor binding overlap with recently identified epitopes of neutralizing antibodies. Firstly, the nicotinic acetylcholine receptor (nAChR)-binding peptide at domain III residues 175-203 [16, 37, 38] falls within the epitopes of mAbs CR57 and RVC20, as discussed in more detail below. Secondly, mAbs 62-7-13 [11], RVA122 [15], 17C7 [16], CR4098 [22, 23], and 523-11[18] bind proximal to putative p75 neurotrophin receptor (p75NTR)-engaging residues K330 and R333 in domain I [39].

### Structure-based analysis indicates that mAb CR57 may interfere with nAchR binding and fusogenic transitions

nAChR, a pentameric transmembrane protein involved in signal transduction at post-synaptic sites in neuromuscular junctions and the central nervous system, was the first receptor described for RABV [37, 40, 41]. The interaction between RABV-G and nAChR is postulated to contribute to the behavioral changes characteristic of rabies disease, given its potential to disrupt cell signaling events orchestrated by certain nAChR subtypes in the central nervous system, altering host behavior towards hyperactivity [38, 42]. Furthermore, studies focused on identifying functional regions in rabies spike found that residues 175-203 share sequence similarity with snake venom curaremimetic neurotoxins, which bind nAChR with high-affinity [37, 41]. This discovery was corroborated by a surface plasmon resonance study, which reported an interaction between a RABV-G peptide (residues 183-196) and an acetylcholine-binding protein derived from *Lymnaea stagnalis* (L-AchBP, a homologue to nAChR) [43] with a dissociation constant (K_D_) in the micromolar range [42].

Given the involvement of nAChR in RABV entry [2, 3], antibodies that sterically hinder nAChR binding could have an inhibitory effect. We mapped the peptide implicated for nAChR binding onto the RABV-G spike in the context of bound CR57, and found that the antibody occludes access to a segment of the receptor binding peptide (Fig 4). Previous competition assays have established that the minimal nAChR binding region comprises RABV-G residues 187-199 (‘TPCDIFTNSRGKR’) [37, 44], and we note that the majority of this sequence falls within the CR57 epitope. Based on our structural analysis, CR57 could significantly hinder binding to nAChR. Notably, this interpretation is supported by a previous study where mAbs 2PV36.74 and 3PV6.39 raised against RABV-G residues 190-203 were able to inhibit the binding of RABV-G to nAChR [45].

**Fig 4.**
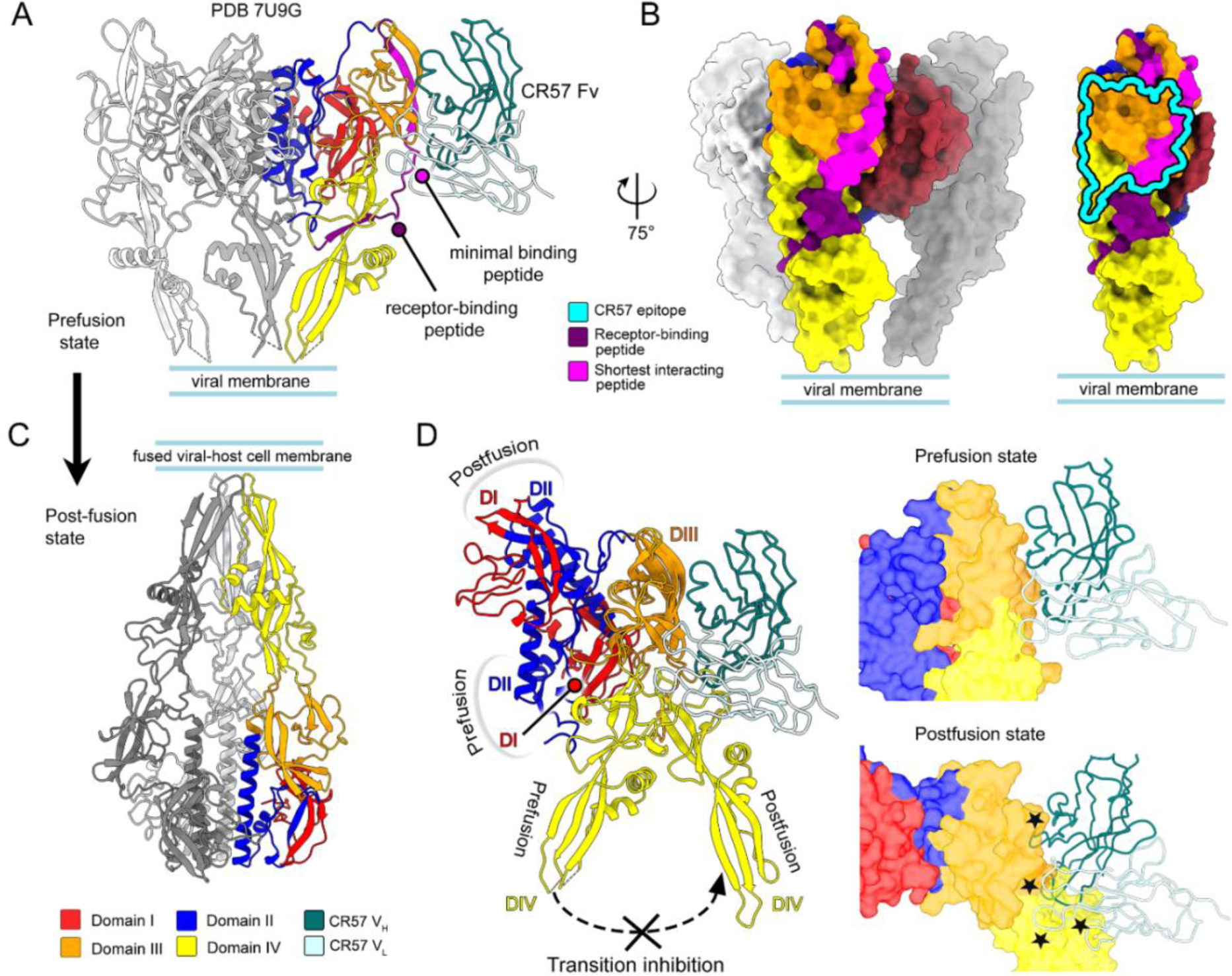
Structural analysis reveals putative mechanisms for CR57-mediated RABV neutralization. (A) Superposition of CR57−RABV-G domain III complex onto the structure of the trimeric prefusion spike (PDB: 7U9G [15], for clarity only a single CR57 Fv is displayed). Panels A and B illustrate the occlusion of the nAChR-binding RABV-G peptide by CR57 Fv. The nAChR binding region is comprised of RABV-G residues 175-203, colored dark magenta, with the minimal binding peptide residues 187-199 highlighted in magenta. (B) Surface representation of the RABV-G spike, with the CR57 epitope outlined by a cyan border, shows that the epitope occludes the half of the minimal binding region for nAChR engagement. (C) AlphaFold2 predicted model of the postfusion RABV-G spike. (D) Superposition of the predicted postfusion RABV-G model and the prefusion RABV-G structure (with CR57 shown bound to domain III) illustrates the steric incompatibility of CR57 Fv with the postfusion state. Steric clashes are denoted by stars.

Another possible mechanism of neutralization is the inhibition of the conformational transitions of the RABV spike from prefusion to postfusion state. Following endocytotic uptake of the viral particle, endosomal acidification triggers spike rearrangement into a postfusion conformation [16, 18, 33], driving membrane fusion that delivers the viral genome into the host cytoplasm. As the postfusion structure of RABV-G remains unknown, we employed AlphaFold2 [46] to acquire a high-confidence structure prediction of the postfusion spike for our analysis of mAb CR57 binding in the postfusion context. We aligned the CR57 diabody−RABV-G domain III structure to the predicted postfusion spike model to assess the effects of conformational changes on the epitope (Fig 4C). While conformational changes in domain III itself are minimal, CR57 targets a hinge region located between domain III and IV that closes upon transition to the postfusion conformation, resulting in clashes between bound CR57 and postfusion RABV-G (Fig 4D). Consequently, we expect that the binding of mAb CR57 to the prefusion spike would obstruct the closing of the hinge region, and thus prevent the adoption of the fusogenic conformation required for viral entry.

In summary, our structural analysis indicates that, in addition to targeting an epitope that overlaps with the putative nAChR receptor binding site, CR57 is likely to interfere with the formation of a functional postfusion conformation, presenting two points where the infection cascade may be halted.

## Discussion

Our study contributes to the understanding of antibody-mediated RABV neutralization. We have resolved an epitope in the RABV-G protein that is targeted by the potent and broadly neutralizing mAb CR57. Crystal structure of the CR57−RABV-G domain III complex demonstrates the significance of the ‘KLCGVL’ peptide within domain III as a crucial region in the antibody-antigen complex interface, which both mAb CR57 and mAb RVC20 bind through a largely hydrophobic interface where aromatic residues play a key role. Based on these similarities between the epitopes of CR57 and RVC20, we expect that this binding mode may be predominant across antigenic site i targeting antibodies. Intriguingly, despite its largely hydrophobic nature, the peptide ‘KLCGVL’ is solvent-exposed in the context of both prefusion and postfusion spike conformations. Moreover, it is well-conserved across lyssavirus phylogroup I [33]. While the exact role of the ’KLCGVL’ peptide remains unknown, we hypothesize that in addition to the role of C228 in the folding of RABV-G, the peptide may play a role in receptor binding or in the intermediate conformational states during membrane fusion, imposing functional constraints on residue variation in this region. We note that another peptide proximal to ‘KLCGVL’, ‘CDIFT’ within domain III (residues 189-193), is also conserved across lyssavirus phylogroup I [33] and confirmed to interact with rabies receptor nAChR [37, 42]. The similar level of conservation observed in both peptides could indicate a shared functionality in receptor engagement. We postulate that mAbs targeting this region, where antibody binding is incompatible with the postfusion conformation, may additionally prevent receptor binding.

This study presents the first instance where a diabody was used to obtain an antibody-antigen complex crystal structure. It should be noted that our attempt to crystallize Fab CR57 in complex with RABV-G domain III did not yield diffracting crystals, highlighting the potential of diabodies as crystallization aids. The identical neutralization behavior of CR57 diabody and Fab indicates that the interface presented by our diabody-domain III complex represents the native mode of antigen engagement. We note that while a diabody has the potential to bind to two antigens simultaneously, which would likely increase neutralization potency, its neutralizing activity remains similar to that of a Fab. We attribute this similarity in neutralizing activity to the low density of spikes on the viral surface.

In summary, RABV-G domain III, with its antigenic and solvent-exposed characteristics, emerges as one of the prime targets for monoclonal antibody-based targeting. By identifying the epitopes of broadly neutralizing antibodies such as mAb CR57, we gain a deeper understanding into how rabies virus is targeted by the humoral immune response, and outline functionally critical regions in the lyssavirus spike.

## Materials and Methods

### Recombinant protein expression and purification

The CR57 diabody was designed based on the reported sequence of CR57 mAb (Genbank AAO17821.1 and AAO17824.1), obtained as codon-optimized synthetic cDNA from GeneArt (Thermo Fisher Scientific), and cloned into the pHL-sec mammalian expression vector (Addgene #99845) [47]. CR57 diabody expression vector was transiently transfected into Expi293F cells at 3×10^6^ cells/ml, using the reagents and protocols of the Expi293^TM^ expression system kit (Gibco^TM^, ThermoFisher Scientific, catalogue no.14635). The cells were cultivated on an orbital shaker at 125 rpm, 37°C with 5% CO2 for six days. CR57 diabody-containing supernatant was collected, clarified by centrifugation and sterile filtered (0.45 μm Corning). Filtrated supernatant was purified by immobilized nickel affinity chromatography with a 1-ml HisTrap excel column (Cytiva) using 300mM imidazole for elution. Finally, the sample was purified by size exclusion chromatography (SEC) using a Superdex 200 10/300 Increase column in an ÄKTA Pure FPLC system (Cytiva), in 10 mM Tris (pH 8.0)–150 mM NaCl buffer.

In order to produce domain III, a construct encoding RABV-G domain III residues 32-59 and 181-259 (GenBank AAC34683.1) connected by a linker comprised of three glycines was designed, commercially obtained as codon-optimized synthetic cDNA from GeneArt (Thermo Fisher Scientific), and cloned into the pURD vector [48]. An Expi293F suspension cell line stably expressing the protein was then generated as described by Seiradake *et al*. [49]. In short, Expi293F cells were plated to a 6-well plate (Greiner Bio-One) and, following overnight cultivation, co-transfected with the domain III/pURD construct and an integrase expression vector pgk-φC31/pCB92 (Addgene #18935) [50]. Following overnight cultivation, the cells were transferred to a T75 flask (Thermo Fisher Scientific), and upon reaching confluence, selected with passaging under 2μg/ml puromycin selection for 10 days. After puromycin selection, cells were detached with trypsin treatment, re-transferred to suspension cell culture in Erlenmeyer shaker flasks (Corning) and upscaled for purification. The supernatant containing domain III was collected and purified via successive nickel affinity chromatography and SEC, as described above for CR57 diabody. CR57 diabody:domain III complex formation was validated by analytic SEC on a Superdex 200 Increase 3.2/300 column (Cytiva), and crystallization samples were produced by mixing purified CR57 diabody and domain III in a 1:1 molar ratio to produce a crystallization sample at 4.8 mg/ml.

### Crystallization and structure determination

CR57 diabody and CR57 diabody:domain III complex were crystallized individually by sitting drop vapor diffusion at 20°C. CR57 diabody was crystallized at 7 mg/ml, in a condition comprising 0.2M magnesium chloride, 0.1M bis-tris, pH 5.5 and 25 w/v polyethylene glycol 3350. The CR57 diabody−domain III complex at 4.8mg/ml was crystallized in a condition containing 0.1M MD Morpheus Buffer System 3, pH 8.5, 0.1M MD Morpheus Carboxylic Acids Mix and 37.5v/v MD Morpheus MPD_P1K_P3350 Mix. The crystals were immersed into a cryo-protectant containing 26% (vol/vol) glycerol followed by rapid transfer into liquid nitrogen. X-ray diffraction data were recorded at beamline I24 (λ = 0.99987 Å), Diamond Light Source synchrotron, United Kingdom.

Data were indexed, integrated, and scaled with XIA2 using the 3dii pipeline [51]. The structure of apo-CR57 diabody and CR57 diabody:domain III complex was solved by molecular replacement in Phenix-MR [52]. Homology model of CR57 Fv was generated using the SWISS-MODEL server [53] based on PDB 5GRX [30] as a search model for the diabody component, whereas PDB 6TOU [33] was used as a search model for domain III in the complex structure along with the Fv model described above. Structure refinement was performed by iterative refinement using Phenix [52] and Coot [54], which was used for manual rebuilding between refinement cycles. MolProbity [55] was used to validate the model. Molecular graphics images were generated using PyMOL (The PyMOL Molecular Graphics System, Version 2.5.0, Schrödinger, LLC) and UCSF ChimeraX [56, 57].

### RABV-G pseudotyped rVSV production

Reporter gene *Renilla* luciferase was cloned into the multiple cloning site MCS-2 of plasmid pVSV-ΔG-N/P-MCS2-2.6 (Kerafast, Boston Massachusetts), which encodes the positive-sense RNA of a replication-defective recombinant vesicular stomatitis virus (rVSV) where the glycoprotein (G) gene has been replaced with a unique multiple cloning site (MCS-1) and an additional transcription start-stop sequence with MCS-2 has been inserted between the nucleoprotein (N) and phosphoprotein (P) genes. The resulting plasmid pVSV-ΔG*RenLuc was used to rescue a replication-defective recombinant *Renilla* luciferase expressing virus (rVSV-ΔG*RenLuc) bearing VSV-G envelope protein as previously described [58].

For pseudotyping rVSV-ΔG*RenLuc with rabies spike protein the residues 1 - 485 including the ecto- and transmembrane domains but excluding the cytoplasmic tail of RABV-G (ERA strain, UniProtKB ID: P03524.1, residue numbering includes signal peptide) were cloned into pEBB vector (Addgene #22226), resulting in plasmid pEBB-R0CT. HEK 293T cells were plated in three 10 cm plates at a seeding density of 2.2×10^6^ and after an overnight incubation at 37⁰C in a 5% CO_2_ incubator, the cells were transfected with 21μg pEBB-R0CT using our *in-house* calcium phosphate transfection reagent according to previously described protocols [58]. At 48 hours post transfection, media was removed from the plates and the cells were infected with 1ml of rVSV-ΔG*RenLuc and left to adsorb for 1 hour at 37⁰C in a 5% CO_2_ incubator. The inoculum was removed, and the plate was washed with PBS, prior to the addition of fresh media. The plate wase then incubated at 37⁰C in a 5% CO_2_ incubator for 48 hours. Supernatant from the plate was harvested, clarified by low-speed centrifugation, titrated, and stored at +4⁰C.

### RABV-G pseudotyped VSV neutralization assay

Neutralization assay was performed on Vero E6 cells (ATCC VERO C1008) that were grown to 100% confluence on a 96-well black Polystyrene Microplate (Corning, New York, USA) with a flat clear bottom. Two-fold serial dilutions (1/400-1/8192000) from CR57 Fab or CR57 diabody, using a 40uM stock concentration for both, were made in U-bottom 96-well microplates (Corning, New York, USA) to a final volume of 30μl. A 1 in 100 dilution of the supernatant containing R0CT pseudotype was chosen via titration as the working dilution since it gave a luminescent count of about 10,000 and the cells exhibited minimal damage due to VSV cytopathic effect. A 30 μl volume of this working R0CT pseudotype dilution was added to the wells, and the mixture was incubated at 37⁰C in a 5% CO_2_ incubator for 1 hour. The mixture was then used to inoculate the previously mentioned confluent Vero E6 plates and incubated under the same conditions for 18 hours. Luminescence in the wells was counted using the *Renilla*-Glo® luciferase assay system (Promega, Madison, Wisconsin) according to the manufacturer’s specifications, read on the HIDEX sense microplate reader with software version 0.5.11.2, counting from the top and using 5 seconds counting time. The percentage neutralization in each well was calculated by comparing the counts in the test wells to those of the virus-only negative control, which were prepared by using media in place of diabody or Fab.

### AlphaFold2 structure prediction

The sequence of RABV-G ectodomain residues 1–439 (numbering does not include the signal peptide; GenBank AAC34683.1) was used to generate a model of the RABV-G postfusion structure using AlphaFold v.2.2.4 [46] on ColabFold v.4.1.4 [59]. Prediction used 266 unique sequences and was performed with six recycles and relaxation enabled. The mapping of each residue’s predicted local-difference test (pLDDT) confidence score on the model is presented in Fig S4.

### Accession Codes

Atomic coordinates and structure factors of the apo-CR57 diabody and CR57 diabody−RABV-G domain III complex have been deposited in the PDB (accession codes XXXX and YYYY, respectively).

## Acknowledgments

Sara Stébe, Virpi Syvälahti and Anna Mäkelä are gratefully acknowledged for their work in generating plasmids for VSV pseudotyping. The facilities and expertise of the HiLIFE Crystallization unit at the University of Helsinki, a member of FINStruct and Biocenter Finland, are gratefully acknowledged. We thank Diamond Light Source for beamtime (proposal MX26794-20) and the staff of beamline I24 at Diamond Light Source for assistance with data collection. Finally, we are thankful to our funders, Academy of Finland (grant # 342988 and # 346508 to I.R.) and the University of Helsinki (3-year grant awarded to I.R.).

