## Supporting Information for "Structural insight into rabies virus neutralization revealed by an engineered antibody scaffold"

### Items:

**Table S1.** Crystallographic data collection and refinement statistics for diabody CR57 alone and in complex with RABV-G domain III.

**Table S2.** An overview of antigenic sites on the RABV-G ectodomain based on existing literature.

**Figure S1**. Conformational differences between the structure of CR57 diabody and the structures of previously reported monospecific diabodies.

**Figure S2**. Comparison of domain III structures and the positioning of CR57 diabody–RABV-G domain III complex in the context of the prefusion spike.

**Figure S3**. LigPlot+ diagram of the interactions at the diabody CR57–RABV-G interface.

**Figure S4**. AlphaFold2 structure prediction of the postfusion RABV-G spike.

Supplementary References

**Table S1.** Crystallographic data collection and refinement statistics for diabody CR57 alone and in complex with RABV-G domain III.

| | Apo-diabody CR57 | Diabody CR57–RABV-G domain III complex |
|---|---|---|
| **Data collection** | | |
| Space group | $P\ 2_1\ 2_1\ 2_1$ | $C$ 1 2 1 |
| Cell dimensions | | |
| $a$, $b$, $c$ (Å) | 64.8, 65.0, 122.3 | 195.7, 57.4, 72.8 |
| $\alpha$, $\beta$, $\gamma$ (°) | 90.0, 90.0, 90.0 | 90.0, 94.3, 90.0 |
| Resolution (Å) | 57.24-2.38 (2.42-2.38)* | 48.81-3.15 (3.20-3.15)* |
| $R_{\text{merge}}$ | 0.23 (1.34) | 0.21 (0.73) |
| $R_{pim}$ | 0.07 (0.50) | 0.12 (0.45) |
| I/σ I | 9.5 (1.7) | 5.7 (1.4) |
| $CC_{1/2}$ | 0.99 (0.49) | 0.96 (0.60) |
| Completeness (%) | 100.0 (100.0) | 99.6 (96.0) |
| Multiplicity | 11.7 (7.9) | 4.1 (3.1) |
| **Refinement** | | |
| Resolution (Å) | 44.5-2.38 (2.42-2.38) | 48.8-3.15 (3.20-3.15) |
| No. reflections | 21356 | 14170 |
| $R_{\text{work}}$ / $R_{\text{free}}$ | 0.19 / 0.23 | 0.25 / 0.29 |
| No. atoms | | |
| Protein | 3601 | 4921 |
| Ligand/ion | - | - |
| Water | 66 | - |
| *B*-factors | | |
| Protein | 65.7 | 103.3 |
| Ligand/ion | - | - |
| Ramachandran plot (%) | | |
| Favored region | 97.47 | 95.44 |
| Allowed region | 2.53 | 4.56 |
| Outliers | 0.00 | 0.00 |
| R.m.s deviations | | |
| Bond lengths (Å) | 0.001 | 0.003 |
| Bond angles (°) | 0.439 | 0.758 |

* Values for the highest resolution shell are shown in parentheses

**Table S2**. An overview of antigenic sites on the RABV-G ectodomain based on existing literature. Data is sourced from references [1-6].

| Site name | Residue range * | Motif | Site location | Antibodies targeting the site |
|---|---|---|---|---|
| Site i | 226-231 | KLCGVL | Domain III | CR57, RVC20, RVC3, 62-71-3, 10EC9, M725, W509-6, M818, 16EH11, W101-1, M1089 |
| Site iia | 198-200 | KRA | Domain III | W110-3 |
| Site iib | 34-42 | GCTNLSGFS | Domain II and Domain III | E559, M727-5-1, M777-16-3, M778, M1094, 1112-1, RV01, RV03, RV05, RV08, RV09 |
| Site iii | 330-338 | KSVRTWNEI | Domain I | CR4098, RAB1 or 17C7, RVA122, 523-11, 62-7-13,<br>10ED8, W1120-10, M1100, RVC58, RVC38, RVA144, RVB492, RVC111, RVC21, RVB185 |
| Site iv | 251 | W | Domain III | 15-13 |
| Minor site 'a' or G1 | 342-343 | KG | Domain I | M724, W120-6, 16AH8, M785 |
| G5 | 261-264 | HDFH | Domain II | AR16, M1078 |

* residue numbering excludes the signal peptide

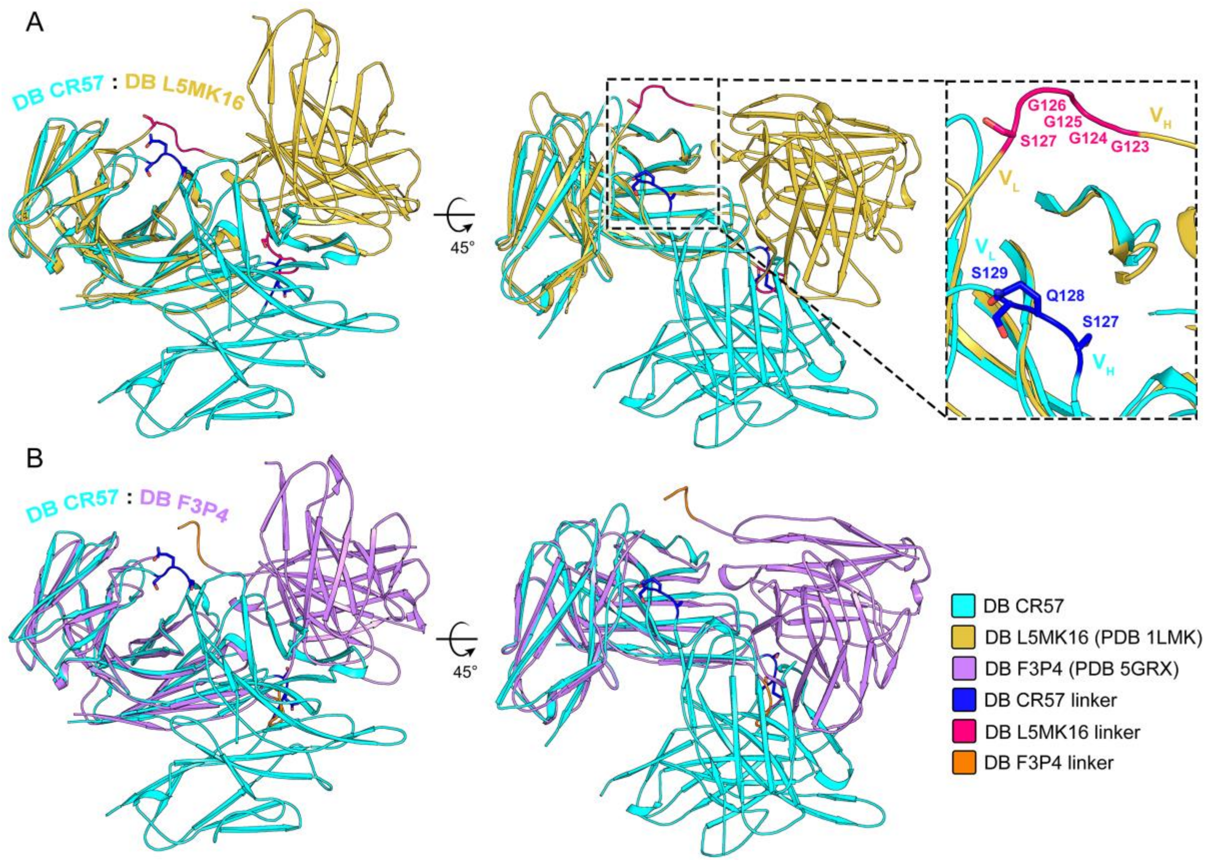


**Fig S1.** Conformational differences between the structure of CR57 diabody and the structures of previously reported monospecific diabodies. (A) Superposition of CR57 diabody (DB), colored cyan, and diabody L5MK16 [7], colored golden orange. The zoom panel shows the linker residues between the $V_H$ and $V_L$ regions. (B) Superposition of CR57 diabody (DB), colored cyan, and diabody diabody F3P4 [8], colored lavender. While individual Fv pairs have a congruent fold across all structures, the orientation of the Fv pairs relative to each other varies.

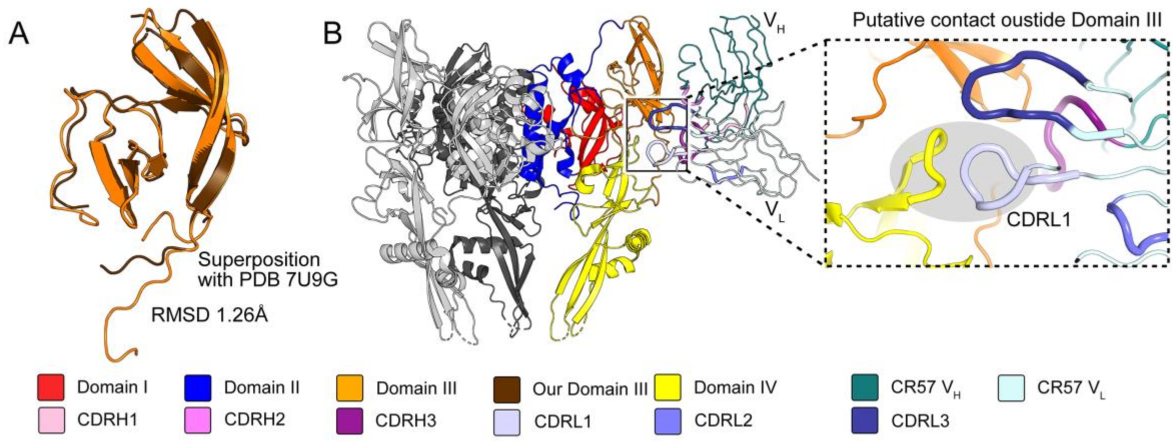


**Fig S2.** Comparison of domain III structures and the positioning of CR57 diabody–RABV-G domain III complex in the context of the prefusion spike. (A) Superposition of our RABV-G domain III structure with the previously reported structure RABV-G domain III from a trimeric prefusion RABV-G spike (PDB 7U9G [9]). (B) Superposition of CR57 diabody–RABV-G domain III complex on the full structure of the trimeric spike ectodomain (for clarity, only a single CR57 Fv region is shown). While domain III comprises the majority of the interface (see main Fig 2 and Fig S3), the superposition indicates that CDRL1 is pointed outside of domain III, and may engage with a region in domain IV in the context of the trimeric prefusion spike.

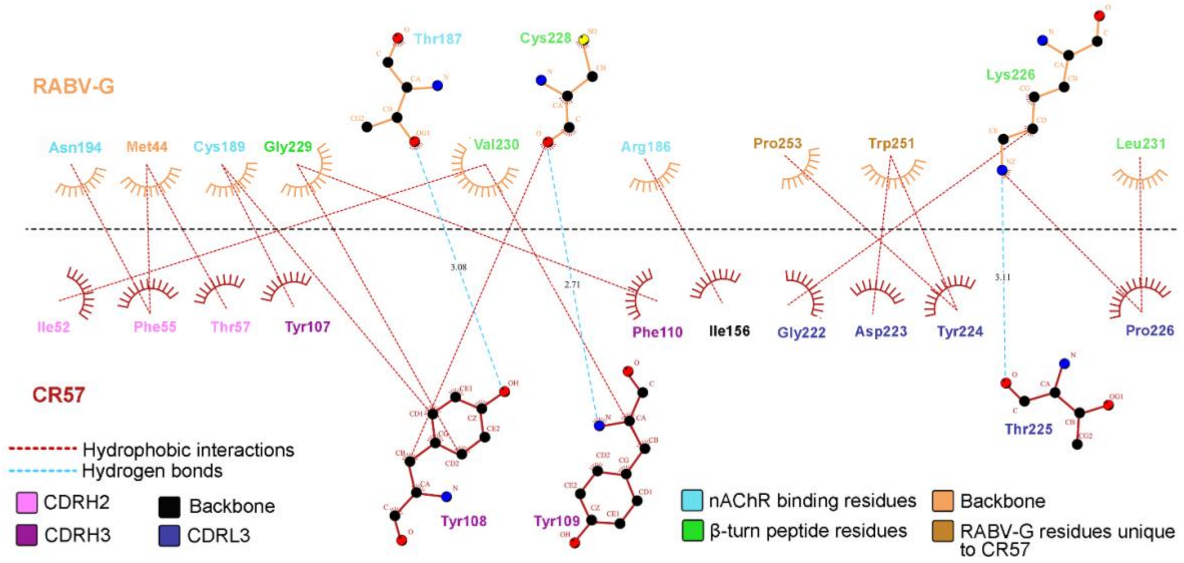


**Fig S3.** LigPlot+ diagram of the interactions at the diabody CR57–RABV-G interface. RABV-G residues are labelled using three different colors: cyan, green, and orange, corresponding to their involvement in nAChR-binding, β-turn peptide-binding, and CR57 interactions, respectively. The dark orange RABV-G residues are unique to the interface when compared to other RABV-G domain III targeting mAbs, such as RVC20 and 1112-1. The CDR residues are labeled in shades of pink and dark blue, representing CDRH2, -H3, and -L3, respectively. Carbon, nitrogen, and oxygen atoms are represented as black, blue, and red spheres, respectively. Residues engaged in hydrogen bonding are depicted as sticks, while those involved in hydrophobic interactions are shown as spoked arcs. Hydrogen bonds are represented by cyan dotted lines, and hydrophobic interactions are depicted by red dotted lines. The diagram was generated using LigPlot+ [10].

.

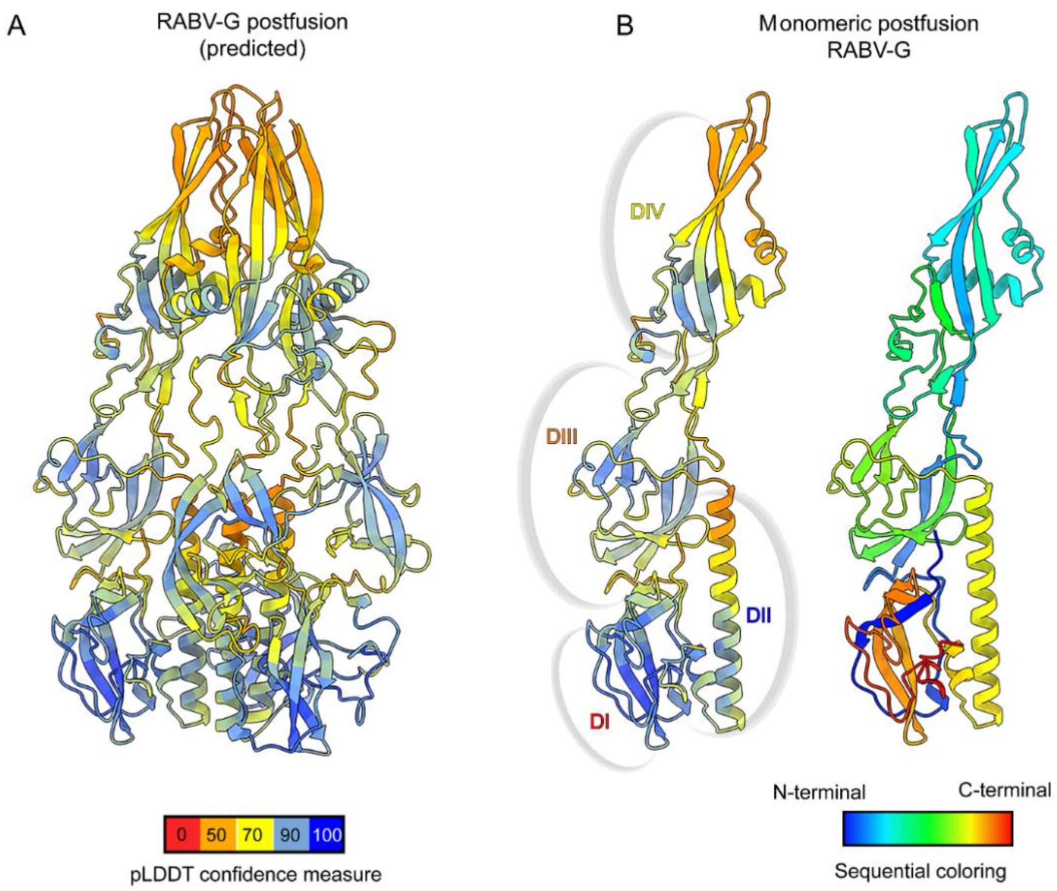


**Fig S4.** AlphaFold2 structure prediction of the postfusion RABV-G spike. (A) An AlphaFold2 [11] model of the postfusion RABV-G spike colored according to the predicted local-difference test [pLDDT] confidence score of each residue. 77% of the modeled 439 residues have a pLDDT score >70, indicating high confidence prediction. (B) Illustrations of a single chain from the predicted posfusion RABV-G structure, colored based on pLDDT confidence scores (right), and the same chain colored rainbow ramped from the N-terminus (blue) to the C-terminus (red).
